# Electron microscopic analysis of the exo-skeleton of hadal zone amphipod *Hirondellea gigas*

**DOI:** 10.1101/448175

**Authors:** Hideki Kobayashi, Hirokazu Shimoshige, Yoshikata Nakajima, Wataru Arai, Hideto Takami

## Abstract

The amphipod *Hirondellea gigas* inhabits the deepest regions of the oceans in extra high-pressure. However, the mechanisms by which they adapt to their high-pressure environments remain unknown. In this study, we investigated elements of the exoskeleton of *H. gigas* captured from the deepest points of the Mariana Trench. The *H. gigas* exoskeleton contained aluminum, as well as a major amount of calcium carbonate. Unlike other accumulated metals, aluminum was distributed on the surface of exoskeletons. To investigate how *H. gigas* obtains aluminum, we conducted a metabolome analysis and found that gluconic acid/gluconolactone was capable of extracting metals from the sediment under the habitat conditions of *H. gigas*. The extracted aluminum ions are transformed into the gel state of aluminum hydroxide in alkaline seawater, and this gel covers the body to protect the amphipod. The aluminum gel would be one of good materials to adapt to such high-pressure environment.

## Introduction

The deepest bottom of the ocean is an extreme environment characterized by extra-high pressures, low temperatures, and oligotrophy, and few animals can adapt to such extreme environments [1-3]. The amphipod *Hirondellea gigas* is a resident of the deepest points of the Mariana Trench (Challenger Deep), the Philippine Trench, the Izu-Ogasawara Trench, and the Japan Trench, where it inhabits depths greater than 8,000 m [4-8]. It has been reported that *H. gigas* produces a number of polysaccharide 2 hydrolases as digestive enzymes and survive in oligotrophic environments by obtaining sugars from degradable plant debris using such enzymes [6, 7]. We attempted to capture animals using baited traps along deep-sea points, and these amphipods were the only catch [5-7]. Amphipods are the main prey for deep-sea fish, and the distribution of deep-sea snailfish slightly overlaps with that of *H. gigas* at approximately 8,000 m [9, 10]. The deep-sea snailfish have been found at depths of 8,145 m in the Mariana Trench, which is the deepest record of fishes [11]. The bottom of the deep trench around 10,000 m in depth seems to be advantageous for *Hirondellea* species to escape from their predators. The extra-high pressure in the deep sea affects the various chemical components of organisms. Calcium carbonate is an important component of crustacean exoskeletons; however, it dissolves in seawater deeper than approximately 4,000-5,000 m (Carbonate Compensation Depth, CCD) [12], and actually they cannot migrate in the deep-sea floor below CCD [13, 14]. Recently, a few species in the crustacean and foraminifera were found in the region slightly deeper than CCD [15-17]. Moreover, lots of foraminifera found in bottom of the Challenger Deep were organic-walled allogromiids, which do not have calcareous wall [18]. Because the habitable zone of *H. gigas* is at depths greater than 8,000 m, they hardly to use calcium carbonate in their exoskeleton. However, the mechanisms by which they adapt to their high-pressure environments remain unknown.

Thus, we investigated elements of the exoskeleton of *H. gigas* specimens captured from the deepest points of the Mariana Trench (Challenger Deep) as well as the Izu-Ogasawara Trench by electron microscopy analyses. We found that aluminum would reinforce calcium carbonate, which is major component of the *H. gigas* exoskeleton.

## Materials and Methods

### Amphipods

The deep-sea amphipod *Hirondellea gigas* was captured from Challenger Deep in the Mariana Trench (depth: 10,898 m) as well as the Izu-Ogasawara Trench (depth: 9,450 m) as described in a previous manuscript [6, 7]. *H. gigas* is >3 cm from the head to tail. We also purchased amphipods from Yokoebi-ya (Fukui, Japan). The coastal amphipods were captured from the seashore of the Maizuru Bay in Japan, and their size is <2-3 mm from the head to tail. All *H. gigas* or the coastal amphipods were stored in storage bags at −80°C without any selection. We selected amphipods for all analysis, randomly.

### Sediment sample

We collected sediment samples from Challenger Deep in the Mariana Trench (11 22.030°N, 142 26.032°E, depth: 10,897 m) using the unmanned remotely operated underwater vehicle “KAIKO” on May 4, 1996. The sediment sample was stored at −80°C.

## Electron microscopy observations and EDS analyses

### Scanning electron microscopy (SEM) and EDS analysis

The freeze-dried amphipod sample was set on the stage of a SEM, which was covered with a silicon plate and carbon tape to avoid EDS signals from the stage. The exoskeleton of the amphipod was observed with a scanning electron microscope (SEM) (SU6600, Hitachi High-Technologies Co., Tokyo, Japan) under accelerating voltages of 20 kV, which have been previously described, and the elementary components were analyzed by energy-dispersive X-ray spectroscopy (EDS) (X-Max^N^, Oxford).

### Scanning transmission electron microscopy (TEM) and EDS analysis

We removed the exoskeleton from *H. gigas* and washed it with DDW and ethanol to avoid inhibition by the oil component during the electron microscope observations. Pieces of *H. gigas* exoskeleton obtained after a milling treatment were placed on a TEM grid (200 mesh Cu Formvar/carbon-coated grid, JEOL) and observed under a TEM (JEM-2100, JEOL) with an accelerating voltage of 200 kV. A scanning TEM (STEM)-EDS analysis was performed at 200 kV with an accumulation time of 60 s.

### Identification of the calcium carbonate using X-ray powder diffraction (XRD) analysis

The exoskeletons of *H. gigas* were removed from individuals with tweezers and dissecting scissors, and washed with methanol and chloroform. After dried the exoskeletons, we cut and crushed them into powder for XRD analysis. The exoskeleton powder was analyzed by X-ray diffractometry (SmartLab, Rigaku), with a Cu radiation source (Kα =1.5418 Å) at 45 kV and 200 mA. The 2θ scan speed, step width, and range was 21.6746 deg/min, 0.02 deg, and 20 to 50 deg. We identified calcium carbonate in the exoskeleton powder through the database (International Centre for Diffraction Data (ICDD)) search of obtained peak positions.

### Extraction of metal ions and gluconic acid/gluconolactone from amphipod

We carefully removed the exoskeleton from amphipod with tweezers and dissecting scissors. Then, we subdivided the exoskeleton in the 0.1N sodium acetate buffer (pH 4.0), and stirred this suspension with a vortex mixer to extract metal ions and gluconolactone/gluconic acid in the exoskeleton. After centrifugation (2,000 x g, 10 min, 4°C), the supernatant was collected, and the precipitated exoskeleton was suspended in the same buffer. Then, we repeated stirring and centrifugation of suspension. Both supernatants were collected, and used for the measurement of metal ions or gluconolactone/gluconic acid in the exoskeleton. The remained body was also subdivided in the 0.1N sodium acetate buffer (pH 4.0), and mashed by BioMasher II (Nippi Inc, Tokyo, Japan) to extract metal ions and gluconic acid/gluconolactone. A small amount of solution, which leaked from amphipod in the process of removing exoskeletons, was also added. After centrifugation (5,000 x g, 10 min, 4°C), the supernatant was divided into two layers of oil and water. Each layer was collected separately. Then, precipitate was suspended in the same buffer and mashed again. After 7 centrifugation (5,000 x g, 10 min, 4°C), each oil and water layer was collected separately. We gathered each layer to the first extraction, and used for measurements.

### Measurement of aluminum, iron, and gluconic acid/gluconolactone content

The aluminum content of the amphipod extract and water were measured using a fluorometric analysis with 8-quinolinol [19]. We suspended samples in DDW, added 0.2 ml of 1% (wt./vol.) 8-quinolonol (Nacalai Tesque, Kyoto, Japan) and 0.2 ml of 2 N CH_3_COONa, and then added DDW to a volume of 5 ml. After mixing well, the aluminum 8-quinolinol complex was extracted with 1 ml of chloroform. The aluminum content was measured by the fluorescent intensity (excitation: 360 nm, emission: 535 nm). An aluminum chloride solution (Wako Pure Chemical Industries, Ltd.) was used as the reference. The iron content of the extracted sediment or soil was measured at an absorbance of 510 nm using Pack Test Fe (Kyouritu Chemical-Check Lab. Co., Tokyo, Japan) based on the reaction of the Fe^2+^ ion and *o*-phenanthroline after reduction. An iron (II) chloride solution was used as a reference. The D-gluconic acid/gluconolactone content of the amphipods was measured using an E-kit “gluconic 8 acid/D-glucono-∂-lactone” (R-Biopharm AG, Damstadt, Germany). Sodium gluconic acid (Wako Pure Chemical Industries, Ltd.) was used as a reference. We calculated total amount of aluminum or gluconic acid/gluconolactone in amphipod from each measured contents and volume of amphipod extracts.

### Effect of enzymatic degradation of gluconolactone/gluconic acid in the extraction of *H. gigas* on the aluminum extraction ability

We crushed and mashed a *H. gigas* individual with BioMasher II (Nippi Inc., Tokyo, Japan). The mashed samples were centrifuged (3,000 x g, 10 min 4°C), and the body fluid sample was collected. The precipitations were washed with 0.2 ml of DDW, and then supernatant was collected and gathered with the body fluid sample after centrifugation (3,000 x g, 10 min 4°C). The gathered sample was filtrated to remove protein using a 3K Amicon Ultra 0.5 ml filter (Merck, Darmstadt, Germany). The pH and volume of filtrated sample was adjusted to 8.0 with 0.1 N NaOH and 0.5 ml, respectively. The 0.1 ml of samples and the same volume of enzyme reaction solution of 0.2 M Tris-HCl buffer containing 20 mM ATP and 5.0 mM NADP were mixed, and then added 1 U of gluconate kinase and 10 U of 6-phosphogluconate hydrogenase 9 (R-Biopharm AG, Darmstadt, Germany). The enzyme reaction was carried out at 25°C for 20 min. The reaction was stopped by filtration with 3K Amicon Ultra 0.5 ml filter. The pH of filtrate was adjusted to around 5.0 by the addition of HCl. The control reaction was also carried out without enzymes. The 0.1 ml of samples were mixed with the sediment of the Mariana Trench (0.1 g-dry weight) in 0.1 M sodium acetate buffer (pH 5.0). We incubated the mixture for 2 h at 4°C, 100 MPa. After centrifugation of mixture (5,000 x g, 10 min, 4°C), the aluminum content of the supernatant was measured.

### High-pressure experiment with salmon roe

The salmon roe were purchased from a fish market. We washed the salmon roe with ice-cold artificial seawater (IWAKI CO., LTD, Tokyo, Japan) 3 times and then soaked them in the ice-cold artificial seawater. We created 16 tubes that each contained 4 salmon roe and artificial seawater, and we then added AlCl_3_ (final conc.: 10 mM) to half of the tubes and adjusted the pH to 8.0 with 5 M NaOH. The tubes were then pressurized at 100 MPa or 0.1 MPa and incubated at 2°C for 1 day. After decompression, we observed the salmon roe and measured the protein concentration of the artificial seawater using the Bradford method [20].

### Cloning of cytochrome oxidase gene from the coastal amphipods

We extracted DNA from 10-20 coastal amphipods using DNAiso (Takara Bio Inc., Kyoto, Japan). The cytochrome oxidase I (COI) gene was amplified from the extracted DNA solution by PCR using the universal primer pair LCO1490 (5′-GGTCAACAAATCATAAAGATATTGG-3′) and HCO2198 (5′-TAAACTTCAGGGTGACCAAAAAATCA-3′) [21]. PCR amplification with a 50-µl reaction volume was performed using the GeneAmp PCR System 9700 (Applied Biosystems, Carlsbad, CA, USA) with EmeraldAmp (Takara Bio Inc., Otsu, Japan) and the buffer supplied with the enzyme. The PCR conditions were as follows: an initial incubation at 96°C for 30 s, 25 cycles of 98°C for 30 s, an incubation at 55°C for 30 s, another incubation at 72°C for 1 min, and a final extension at 72°C for 5 min. The PCR products were cloned in pT7blue-2 vector (Merck Millipore, MA, USA) and then transformed into *Escherichia coli* DH5a for blue/white selection. The cloned COI gene was amplified from the white colony by PCR using the same primers and conditions. The PCR products were analyzed by electrophoresis on a 1% agarose gel. The gel was purified using Exo-SAP digestion with Exonuclease I (USB Corp., Cleveland, OH, USA) and shrimp alkaline phosphatase (SAP) (Promega, Fitchburg, WI, USA) at 37°C for 20 min and then treated at 80°C for 30 min to inactivate the enzymes. The PCR products were sequenced using the primers described above and DYEnamic ET Dye Terminator reagent (GE Healthcare Life Sciences, Piscataway, NJ, USA) on a MegaBACE 1000 (Amersham Biosciences, Piscataway, NJ, USA) automatic sequencer. The nucleotide sequences were trimmed, assembled, and translated using Sequencher 3.7 software (Gene Codes Corp., Ann Arbor, MI, USA).

### Phylogenetic analysis of the coastal amphipod

A preliminary phylogenetic affiliation for each sequence was determined by conducting a BLAST search. The most closely related sequences with representative Lysianassoidean species sequences and certain outgroup sequences were aligned with our sequences using CLUSTALX, and ambiguous regions were excluded from the alignment. Phylogenetic trees were calculated with the PAML algorithm implemented in the TOPALi package ver. 2.5 [22]. The statistical robustness of the analysis was estimated by bootstrapping with 250 replicates.

### Metabolome analysis of *H. gigas*

A metabolome analysis was conducted from 1 frozen *H. gigas* individual using capillary electrophoresis time-of-flight mass spectrometry (CE-TOFMS) and liquid chromatograph time-of-flight mass spectrometry (LC-TOFMS). The metabolome measurements were conducted through the services of a facility at the Human Metabolome Technology Inc., Tsuruoka, Japan.

### CE-TOFMS

Approximately 45 mg of a frozen individual was plunged into 1.5 ml of 50% acetonitrile/Milli-Q water containing internal standards (H3304-1002, Human Metabolome Technologies, Inc., Tsuruoka, Japan) at 0°C to inactivate the enzymes. The individual was homogenized three times at 1,500 rpm for 120 s using a tissue homogenizer (Shake Master neo, Bio Medical Science, Tokyo, Japan), and then the homogenate was centrifuged at 2,300 × g at 4°C for 5 min. Subsequently, 800 µL of the upper aqueous layer was centrifugally filtered through a Millipore 5 kDa cutoff filter at 9,100 × *g* at 4°C for 120 min to remove proteins. The filtrate was centrifugally concentrated and re-suspended in 50 µl of Milli-Q water for the CE-MS analysis. A CE-TOFMS analysis was conducted using an Agilent CE Capillary Electrophoresis System equipped with an Agilent 6210 TOF mass spectrometer, Agilent 1100 Isocratic HPLC pump, Agilent G1603A CE-MS adapter kit, and Agilent G1607A CE-ESI-MS sprayer kit (Agilent Technologies, Waldbronn, Germany). The systems were controlled by the software Agilent G2201AA ChemStation version B.03.01 for CE (Agilent Technologies). The metabolites were analyzed using a fused-silica capillary (50 μm *i.d.* × 80 cm total length) and a commercial electrophoresis buffer (Solution ID: H3301-1001 for the cation analysis and H3302-1021 for the anion analysis, Human Metabolome Technologies) as the electrolyte. The sample was injected at a pressure of 50 mbar for 10 sec (approximately 10 nl) for the cation analysis and 25 sec (approximately 25 nl) for the anion analysis. The spectrometer was scanned from *m/z* 50 to 1,000. Other conditions were applied as previously described [23-25].

The automatic integration software MasterHands (Keio University, Tsuruoka, Japan) was used to obtain peak information, including the *m/z,* migration time for the CE-TOFMS measurement (MT) and the peak area [26]. Signal peaks corresponding to isotopomers, adduct ions, and other product ions of known metabolites were excluded, and the remaining peaks were annotated with putative metabolites from the HMT metabolite database based on their MTs and m/z values as determined by the TOFMS analysis. The tolerance range for the peak annotation was configured at ±0.5 min for MT and ±10 ppm for *m/z.* In addition, the peak areas were normalized against those of the internal standards, and the resultant relative area values were further normalized by the sample amount.

A hierarchical cluster analysis (HCA) and principal component analysis (PCA) were performed using our proprietary software PeakStat and SampleStat, respectively. The detected metabolites were plotted on metabolic pathway maps using VANTED (Visualization and Analysis of Networks containing Experimental Data) software [27].

### LC-TOFMS

Approximately 45 mg of a frozen specimen was plunged into 0.5 ml of acetonitrile containing 1% formic acid and internal standards (H3304-1002, Human Metabolome Technologies, Inc., Tsuruoka, Japan) at 0°C to inactivate the enzymes. The specimen was homogenized three times at 1,500 rpm for 120 s using a tissue homogenizer (Shake Master neo), and then 167 µl of Milli-Q water was added and homogenized once at 1,500 rpm for 120 s. The homogenate was centrifuged at 5,000 × *g* at 4°C for 5 min. The supernatant was used as a sample for the LC-TOFMS analysis, and a precipitate was homogenized once with 0.667 ml in the same solution at 1,500 rpm for 120 s. The homogenate of the precipitate was centrifuged at 5,000 × *g* at 4°C for 5 min. Both supernatants were mixed and centrifugally filtered through a Nanosep 3K (PALL Co., NY, US) at 9,100 × g at 4°C for 120 min to remove proteins. Solid phase extraction was conducted on the filtrate to remove phospholipids. The filtrate was centrifugally concentrated and re-suspended in 100 µl of 50% (v/v) isopropanol solution for the LC-TOFMS analysis.

The LC-TOFMS analysis was conducted using an Agilent 1200 series RRLC system SL (Agilent Technologies, CA, USA) and an ODS column (2×50 mm, 2 µm) equipped with an Agilent LC/MSD TOF system (Agilent Technologies). The LC analysis was performed using a mobile phase of solution A (H_2_O/0.1%HCOOH) and solution B (isopropanol: acetonitrile: H_2_O (65: 30: 5)/0.1% HCOOH, 2 mM HCOONH_4_) at a gradient of 0-0.5 min: B 1%, 0.5-13.5 min: B 1-100%, 13.5-20 min: B 100%. Negative and positive modes were performed for the cationic and anionic metabolites. The MS system, measurement conditions, and analyses were conducted using the same procedures as described above for the CE-TOFMS analysis.

## Result

### Scanning electron microscopy (SEM)/energy dispersive X-ray spectrometry (EDS) analysis of *H. gigas* exoskeleton

First, we analyzed the elements of the exoskeletons of *H. gigas* captured from the Challenger Deep in the Mariana Trench using SEM/EDS. Our results showed a peak of aluminum as well as other major metals, such as calcium, magnesium, potassium, and sodium, in various parts of the exoskeleton (Fig. 1, Table 1). Calcium was most abundant in metal ions. Aluminum was widely distributed throughout the entire body, and an especially high content of aluminum was observed in the tail (telson) and along the edge the feet (pereopods, or uropods) (Fig. 1b-e, Supplementary Fig. 1). However, silicon, which is the major component of the clay mineral aluminosilicate [28, 29], was not observed; therefore, the aluminum in the exoskeleton was not derived from the aluminosilicate itself in sediment. We repeated the same analysis with other *H. gigas* individuals and obtained similar results, including a high aluminum content in their telson (S1, 2 Fig.). Similar to the amphipods from the Mariana Trench, the *H. gigas* specimens captured from the Izu-Ogasawara Trench also had aluminum in their exoskeleton (S3 Fig.). To identify whether the aluminum accumulated into exoskeleton or adhered to surface of exoskeleton, we observed aluminum existence after washing the surface of *H. gigas* individuals with distilled deionized water (DDW). The aluminum was clearly removed from exoskeletons (Fig. 2). Therefore, the aluminum covered the surface of exoskeleton rather then accumulation in the internal part of exoskeleton. A comparison was performed to investigate the aluminum in the exoskeleton of a shallow-sea, coastal amphipod that was captured from Maizuru Bay in Japan and identified as *Prontogenesia* sp. based on the amino acid sequence of cytochrome oxidase (S4 Fig.). Compared with the deep-sea amphipods, the SEM/EDS analysis of the exoskeletons of *Prontogenesia* sp. did not show an aluminum peak (S5 Fig.). To date, aluminum content has not been reported in the exoskeletons of crustaceans; however, we contend that exoskeletons containing aluminum is a unique property of hadal amphipods.

**Figure 1.**
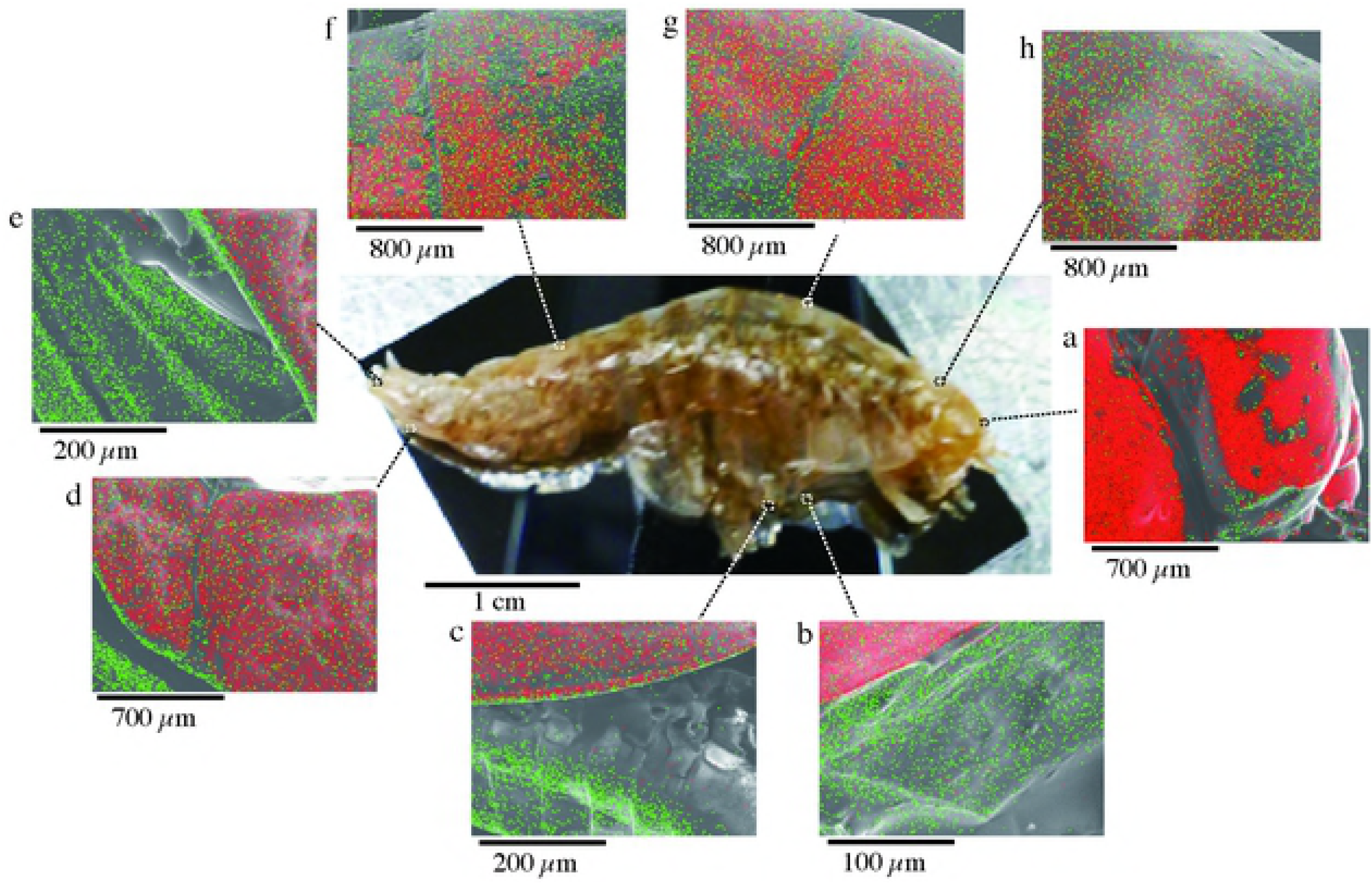
SEM/EDS analysis of the exoskeleton of *H. gigas*. *H. gigas* specimens captured from the Challenger Deep were freeze-dried for the SEM observations. The SEM observations were conducted without any coating. Calcium (red) and aluminum (green) were mapped on the SEM pictures (A, a-i). The EDS spectrum included an annotation of each element with its Kα energy levels (C: 0.284, O: 0.532, Na: 1.071, Mg: 1.253, Al: 1.486, P: 2.013, S: 2.307, Cl: 2.621, Ca: 3.69 (k eV)).

**Figure 2.**
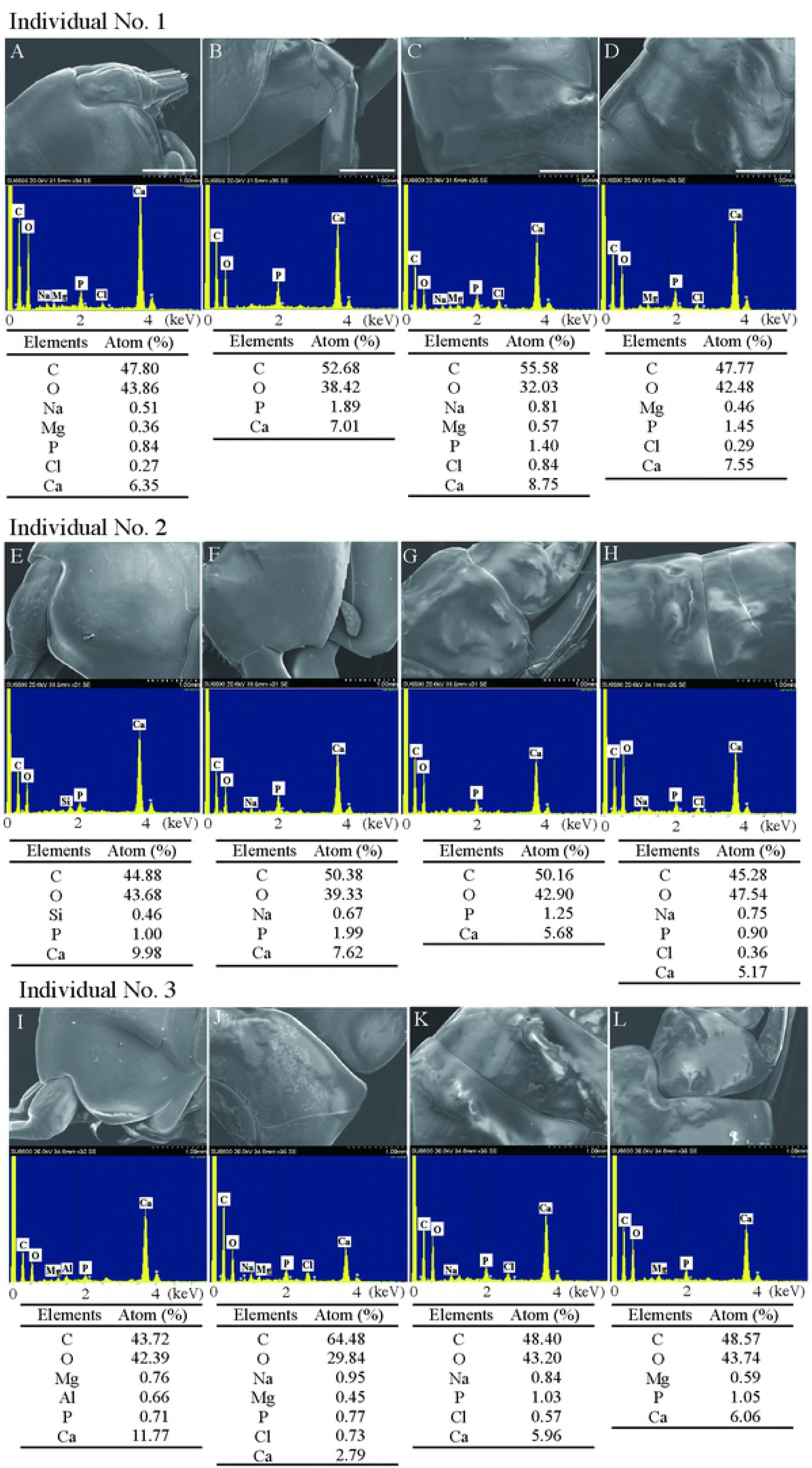
SEM/EDS analysis of the exoskeleton of *H. gigas* after washing with DDW. Three *H. gigas* specimens captured from the Challenger Deep were washed 3 times with DDW, and freeze-dried the SEM observations. The SEM observations were conducted without any coating. Exoskeleton parts of the head (A, E, I), the body (B, F, J), the back (C, G, K), and the telson (D, H, L) were observed. Each panel showed SEM image (top), EDS spectrum (middle), and element composition obtained from EDS spectrum (bottom).

**Table 1.**
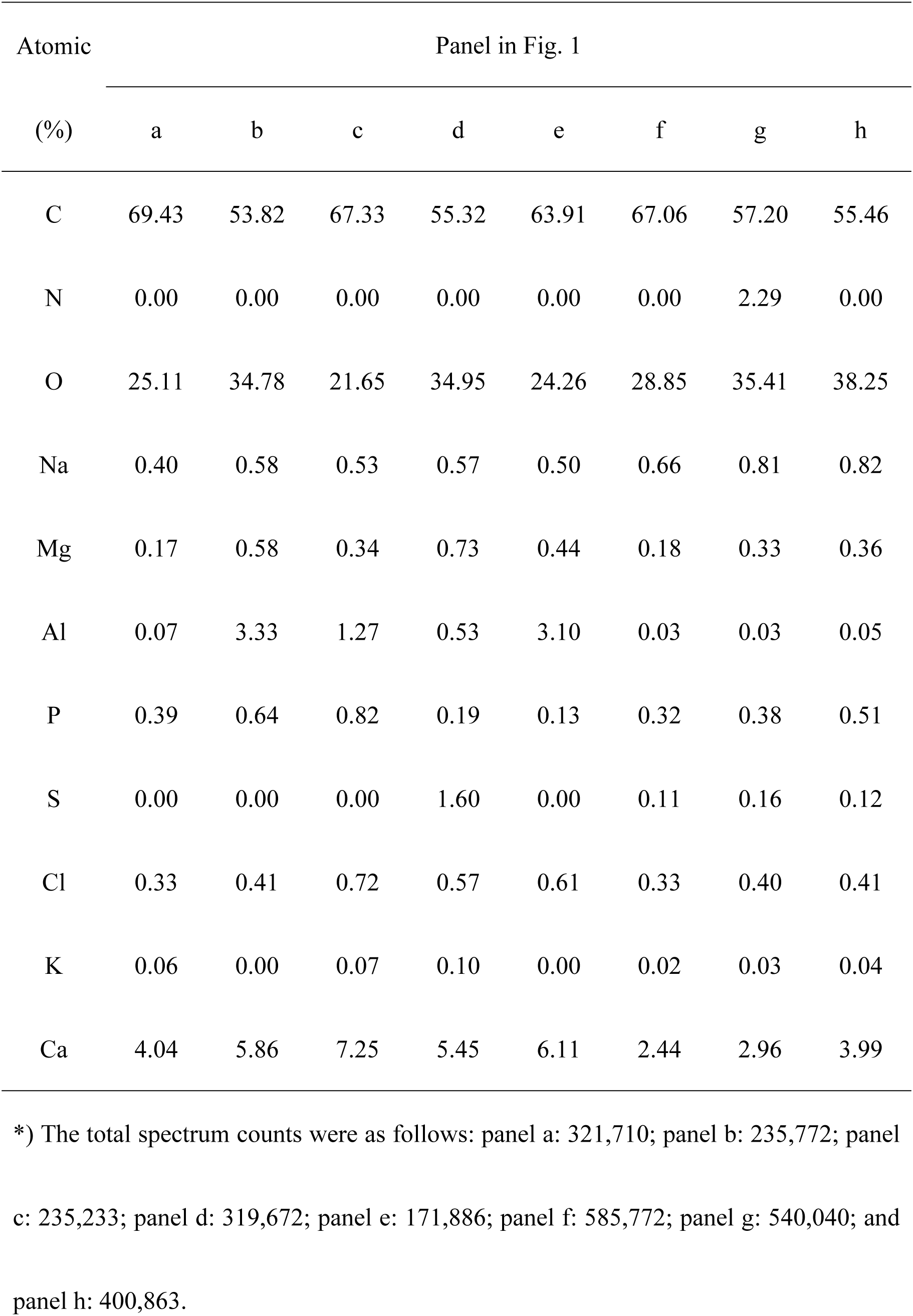
The ratio of atoms was calculated from the total spectrum count* in each panel in Fig. 1

### Scanning transmission electron microscopy (STEM)/EDS analysis of *H. gigas* exoskeleton

We also observed the crushed exoskeleton to find aluminum of the internal exoskeletons of *H. gigas* captured from the Challenger Deep through a scanning transmission electron microscopy (STEM)/EDS analysis (Fig. 3). We found calcium in all crushed exoskeleton, and nickel or copper in a few piece of exoskeleton. Copper and nickel are minor elements in the marine sediment, and *H. gigas* would not accumulate sufficient amounts for detection in the SEM/EDS analysis. The aluminum was not contained in the internal exoskeleton. Silicate and molybdenum were background signals from sample holder. We also observed the internal exoskeletons of the amphipods captured from the Izu-Ogasawara Trench, and did not found any aluminum peak in STEM/EDS analysis (S6 Figure).

**Figure 3.**
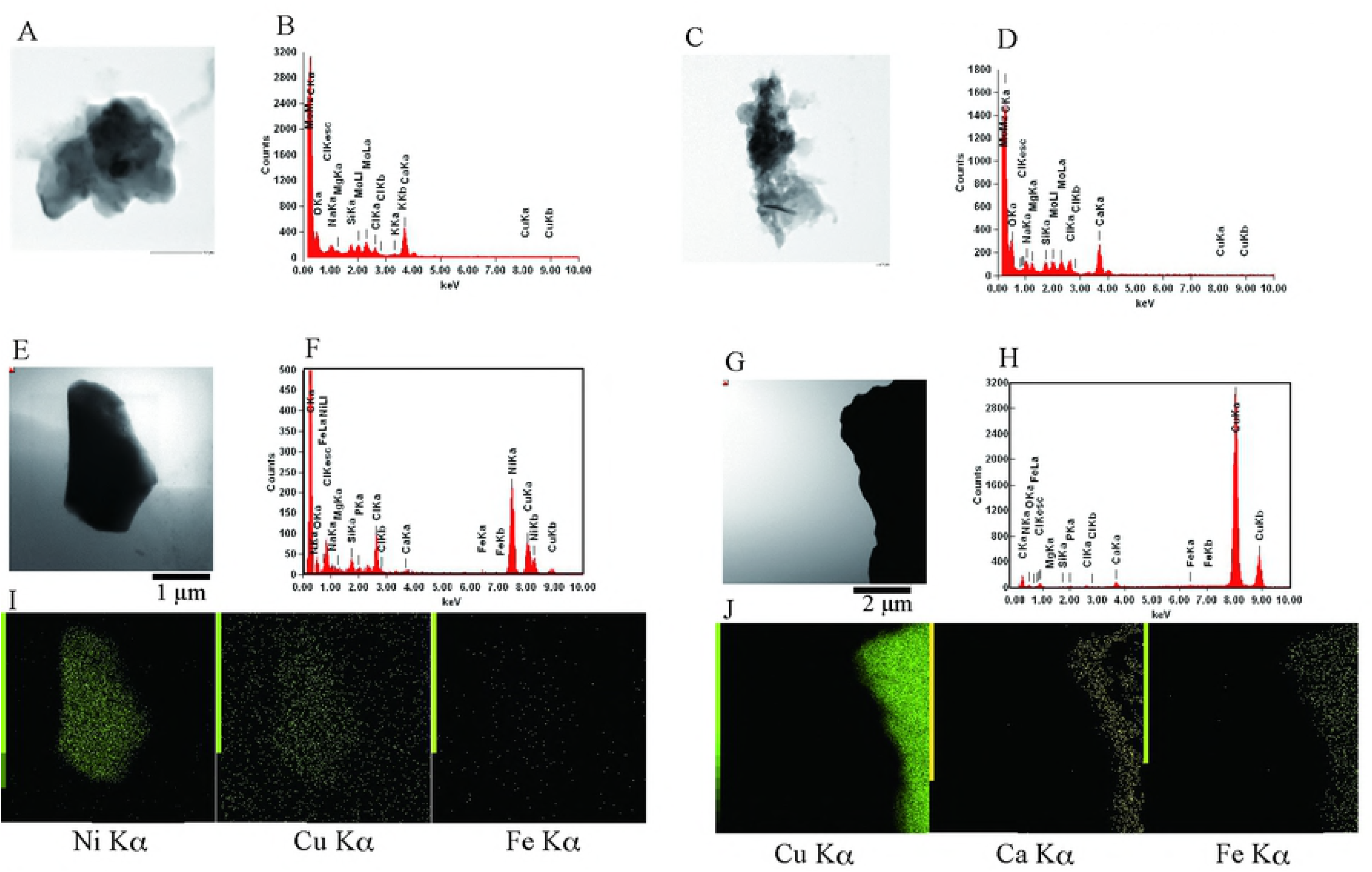
STEM/EDS analysis of pieces of *H. gigas* exoskeleton. Exoskeletons of *H. gigas* captured from the Challenger Deep were removed from the individuals, freeze dried, and then scrapped. Bright-field STEM observations were conducted for pieces of the exoskeletons (A, C, E, G). Characteristic X-rays were collected over 60 s (B, D, F, H), and the major metal signals of panel E and G were mapped (I, J). The Cu or Mo signals were caused by the TEM grid. The Si signal was background.

### X-ray powder diffraction (XRD) analysis of the exoskeletons of *H. gigas*

We found calcium was major metal in the exoskeletons of *H. gigas* through EDS analysis. Generally, it is known that calcium occurs as calcium carbonate and calcium phosphate in the crustacean [30-32], however, much lower peak of phosphorus compared with that of calcium was detected in the exoskeletons of *H. gigas* whose habitat is much deeper than CCD (Table 1). This result indicates the possibility of the existence of calcium carbonate in the exoskeleton. Thus, we carried out XRD analysis of the exoskeletons of 5 individuals in order to examine existence of calcium carbonate (Fig. 4). The diffraction peak positions in 5 samples were the same except for one unknown peak. The 2θ of 7 main peaks was 23.1±0.1, 29.5, 36.0, 39.5, 43.2, 47.7, and 48.6, respectively. Since the crystal material in the exoskeleton was suggested to be trigonal calcium carbonate from database search, calcium detected by the EDS analyses was found to be mainly calcium carbonate. Compared with reference peak of crystal calcium carbonate (CCC), the heights of peaks were decreased in accordance with the increase of 2θ. Because CCC in the exoskeleton was made from amorphous calcium carbonate (ACC), which was synthesized on chitin in crustacean [33-35], the observed peaks are originated from both CCC and ACC. One unknown peak (2θ = 44.6) was found in one sample, and suggested as AlO(OH) (1 1 1) (panel d). However, since we could not find largest peak of AlO(OH) (1 1 0) (2θ = 22.3), we did not conclude that this peak was originated from AlO(OH).

**Figure 4.**
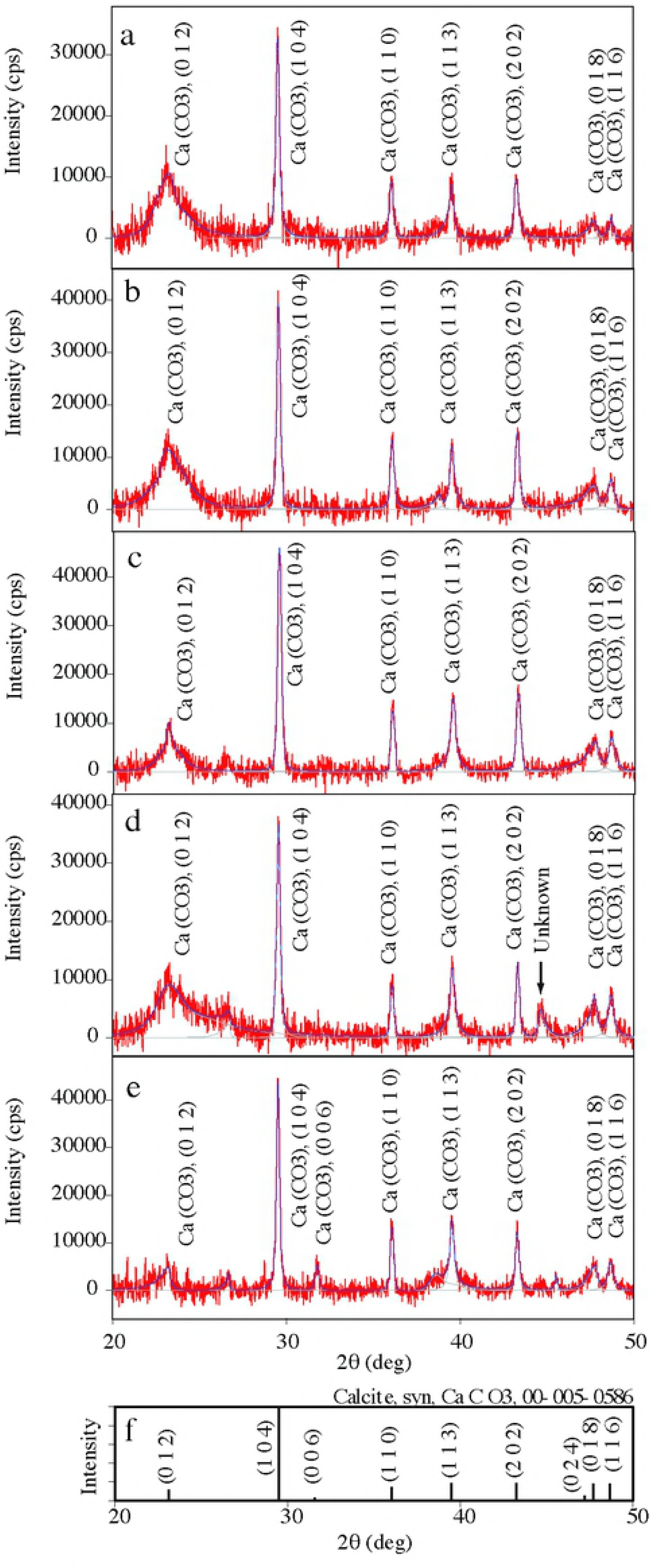
XRD analysis of *H. gigas* exoskeleton. Exoskeleton samples prepared from 5 individuals of *H. gigas* were used for XRD analysis as described in Method (panel A-E). The annotations of peaks were results of database search (panel F). Arrow in panel d indicates unknown peak, which was suggested as AlO(OH) from library search. We cannot identified it as AlO(OH), because other minor peaks were not found.

### Aluminum content in *H. gigas*

Next, we measured the content of aluminum in the *H. gigas* individuals. To avoid the influence of aluminum oxide originating from the sediment, we used the 8-quinolinol method and applied the fluorescent label of the 8-quinolinol-aluminum ion complex [19]. We selected three individuals captured from the Challenger Deep in the Mariana Trench (S7 Fig.). The *H. gigas* exoskeletons of the three individuals contained over 50% aluminum and exhibited only minor differences in content (S1 Table), whereas the body fluid presented varying aluminum content among the 3 individuals. Because the crustaceans inhabiting contaminated areas accumulate metals in their gut [36, 37], the presence of aluminum in the exoskeleton indicated not to be a result of simple accumulation.

### Identification of aluminum extraction agent H. gigas

Because aluminum is the third-most abundant element on Earth, possible origins of the aluminum in *H. gigas* include the marine sediment or the seawater. A large amount of aluminum occurs as aluminum oxide in the clay minerals of the marine sediment [28, 29, 38]. However, aluminum ions have been found in deep-sea water in small amounts (approximately 2 nM) in the North Pacific [39, 40]. When deep-sea amphipods were caught in the baited traps, the content in the digestive organ was the used bait, the sediment, or nothing [5, 6]. We expected that *H. gigas* extracted aluminum from sediment. In fact, an SEM/EDS analysis of another *H. gigas* individual showed the presence of aluminum and silica in the head part, and element mapping showed the same distribution of aluminum and silicon in the head (S8 Fig.). The atomic percent calculated from each peaks were C: 49.2, N: 14.1, O: 30.5, Na: 0.42, Mg: 0.25, Al: 0.47, Si: 1.66, P: 0.22, Cl: 0.32, Ca: 5.1, respectively. The ratio of aluminum and silicon was approximately 1:3.53, which was similar to the aluminum ratio in the sediment (1:3.34-1:4.96) (S2 Table) [41]. Accordingly, the aluminum found in *H. gigas* is likely dependent on the release of aluminum from the marine sediment under the acidic conditions of the gut, whose pH was estimated around 5-6 from the digestive enzyme activities [6, 7, 42]. However, only a limited amount of aluminum was released from the sediment of the Challenger Deep under the same physical conditions as observed in the *H. gigas* gut or habitat (2°C, 100 MPa, and pH 5.0-8.0) [6, 7]. To examine the extraction agent of aluminum from the sediment, we performed aluminum extraction using the protein and non-protein fractions of crushed H. gigas, which are the enzymatic or chemical reactions related to aluminum extraction, respectively. The results showed that the non-protein fraction could extract aluminum (Fig. 5); therefore, we conducted a metabolome analysis of an entire *H. gigas* individual to identify the chemicals that contribute to releasing aluminum from the sediment. The results indicated that *H. gigas* produces 217 chemicals (S3 Table). As potential candidates, we focused on 60 chemicals that are not involved in the metabolic pathways of the animal because metabolites involved in these pathways usually occur in the inner cells and cannot react to extracellular marine sediment in the gut. Among the 60 chemicals, gluconic acid/gluconolactone (gluconolactone in acidic pH) is known as a strong organic chelating agent [43]. A high glucose content (0.43%±0.1% (w/w) (dry weight)) was present in the *H. gigas* individuals, which is a source of gluconic acid/gluconolactone [6]. Furthermore, gluconic acid/gluconolactone levels of 0.3-0.4 mM were measured in the bodies of the *H. gigas* presented here (S4 Table). We examined the effects of gluconic acid/gluconolactone on the release of aluminum from the sediment of the Challenger Deep and confirmed that gluconic acid/gluconolactone released the aluminum ion from the sediment at pH 6.2 or lower under *in situ* conditions (Fig. 6). Under acidic conditions, gluconolactone is a main component in the chemical equation of gluconic acid/gluconolactone and acts as an extraction agent of aluminum from sediment in the gut [43]. Therefore, the extraction of aluminum is a unique property of gluconolactone, rather than a chelating activity of gluconic acid. When we removed gluconic acid/gluconolactone from the extraction of *H. gigas* with enzymatic reaction of gluconokinase (EC 2.7.1.12), the ability of aluminum extraction form sediments was lost (S9 Fig.). Gluconolactone was main aluminum extractor in *H. gigas*. Gluconic acid/gluconolactone can extract minor amounts of iron from sediment under the same deep-sea bottom conditions (S10 Fig.). Therefore, copper and nickel, found in STEM/EDS observation, were also included in and extracted from the sediment by gluconic acid/gluconolactone [29, 43].

**Figure 5.**
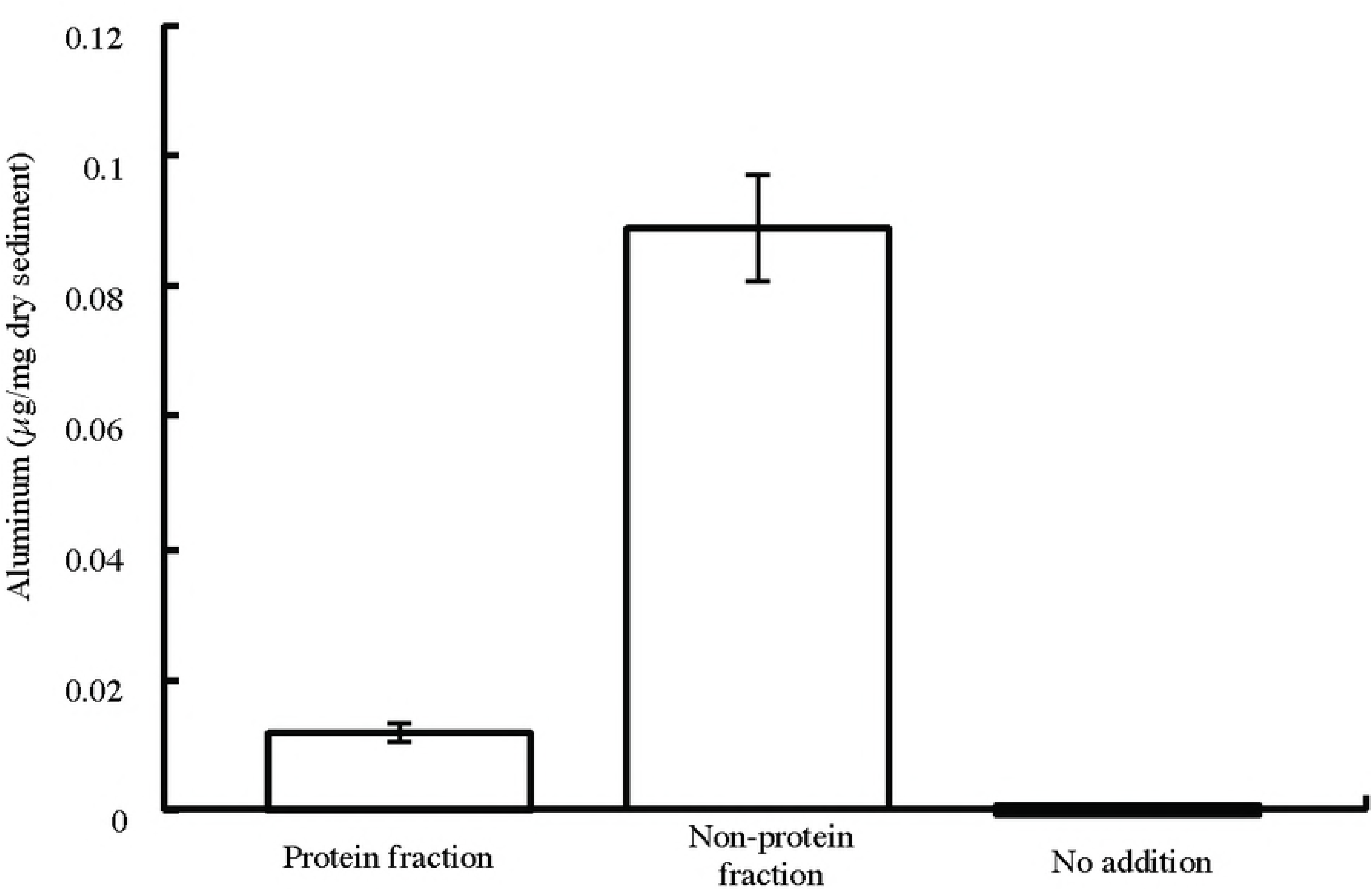
Extraction of aluminum from the sediment of Challenger Deep by the *H. gigas* protein fraction or non-protein fraction. Three *H. gigas* individuals were scrapped and then suspended in 1 ml of (NH_4_)_2_SO_4_ solution at 80% saturation. After incubation at 4°C for 2 h, the *H. gigas* suspensions were centrifuged at 20,000 x *g* for 30 min. The supernatants were used as the non-protein fractions, and the precipitates were suspended in 1 ml of DDW to prepare the protein fractions. We mixed 300 µl of each fraction with the sediment suspension and added sodium acetate buffer to a final volume of 1.8 ml (final concentration: 50 mM, pH 5.0). The mixtures were then pressurized at 100 MPa and incubated at 2°C. After 1 h incubation, the mixtures were de-pressurized and centrifuged at 20,000 x *g* for 10 min. The aluminum content of the supernatants was measured as described in the Methods section. The error bar shows the S.D. (n=3).

**Figure 6.**
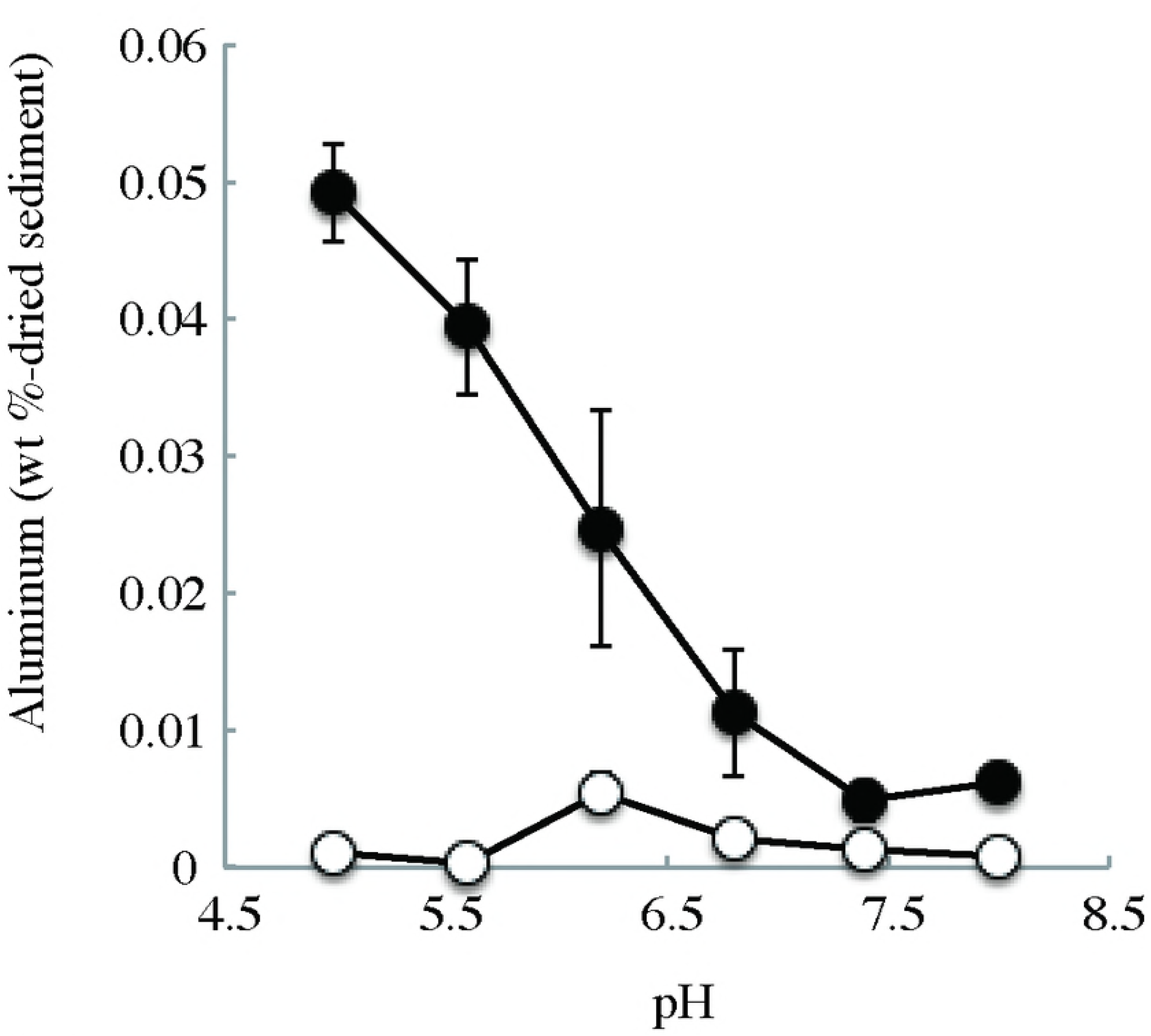
Extraction of aluminum from the sediment of Challenger Deep by gluconic acid/gluconolactone. Sediment samples were washed with distilled deionized water five times and then suspended in each buffer, which contained 10 mM sodium gluconic acid/gluconolactone (closed circle) or not (open circle). Extracted aluminum was measured as described in the Materials and Methods.

## Discussion

We have shown that aluminum hydroxide gel covers the body of *H. gigas* in this study. *H. gigas* inhabits the bottom of the deepest trench by obtaining glucose, a resource of gluconolactone/gluconic acid, from plant debris buried in sediment digested by their own cellulase and hemicellulose hydrolases [5-7]. Namely, *H. gigas* has ability to take in both resources of aluminum and it’s extraction agent from the sediment. Aluminum hydroxide gel is constructed by chemical behavior of aluminum ion toward pH. The extracted aluminum ions in the gut are released to the seawater of around pH 8, where they are transformed to the gel state of aluminum hydroxide and then mainly stored in the *H. gigas* telson [44, 45]. The aluminum spread on the surface of exoskeleton without discharging together with excrement. Thus, it is thought that *H. gigas* has some transport system of aluminum. We also detected gluconic acid in the exoskeletons at 0.06-0.16 µmol per individual (S4 Table), which turns into gluconic acid from gluconolactone in contact with the alkaline seawater.as excrement. Hence, there would be some transporter of aluminum in *H. gigas*. The converted gluconic acid would work as a chelating agent for transferring aluminum ions to whole exoskeleton. The binding property and stability of aluminum hydroxide gel to organisms are still unclear. The aluminum hydroxide gel can be stable only in alkaline environment. Therefore, “aluminum cover” would present only in the lives inhabiting the ocean and some alkaline lakes.

Because the amphipods captured from the Izu-Ogasawara Trench were also found to have the aluminum hydroxide gel, the aluminum hydroxide gel may have some common role for the protection from the high pressure in the deep sea. Thus, we carried out a preliminary experiment to figure out protection effect of aluminum hydroxide gel on high pressure stress using salmon roes, and found less protein leak and no color change in the presence of aluminum hydroxide gel even under 100 MPa (S11 Fig.). The aluminum hydroxide gel may contribute to the adaptation system in the high pressure. However, since the surface of salmon roe is very different from that of amphipod’s exoskeleton, we should do the same experiments using good analogous animal samples of *H. gigas* although it is difficult to find them for better understanding of the properties of aluminum in *H. gigas.*

Discovery of CCC in the exoskeleton of *H. gigas* indicated that CCD did not govern synthesis of biological calcium carbonate. But in contrast to our discovery, the foraminifera living under CCD don’t have calcium carbonate shell [18, 46]. Thus, the difference of structural or chemical composition between amphipod and foraminifera would be related with the retention of calcium carbonate. In any case, the effect of aluminum gel on synthesis and maintenance of crystal carbonate is still unknown.

## Author Contributions

H. K. designed and performed all experiments and wrote the paper. H. K., H. S. and Y. N. prepared the amphipod samples for electron microscopy observations and performed the SEM/EDS and STEM/EDS analyses. H. K., W. A., and H. T. prepared the baited traps and captured *H. gigas* from the Mariana Trench and the Izu-Ogasawara Trench. All authors discussed the results and commented on the manuscript.

## Material and Correspondance

All materials and correspondence should be addressed to Hideki Kobayashi

## Data availability

Sequencing data of COI have been deposited in DDBJ with the accession codes “LC085650-LC085655” (http://getentry.ddbj.nig.ac.jp/getentry/na/LC085650-LC085655).

## Supporting information

**S1 Figure SEM/EDS analysis of the exoskeleton of *H. gigas* telson.**

*H. gigas* specimens captured from Challenger Deep was freeze dried for SEM observations (A, B). Panel B shows an enlargement of the red square in panel A. SEM observations and EDS analyses were conducted without any coating. The EDS spectrum of panel B includes an annotation of each element with its Kα energy level (C: 0.284, O: 0.532, Na: 1.071, Mg: 1.253, Al: 1.486, P: 2.013, S: 2.307, Cl: 2.621, Ca: 3.69 (k eV)) (C). The total spectrum counts were 117,388 in the EDS analysis, and the major elements were mapped (D).

**S2 Figure SEM/EDS analysis of the *H. gigas* exoskeleton.**

An SEM/EDS analysis was conducted on the telson region (A) and the exoskeleton (E) as described in the Methods section. The EDS spectra of panel A and E include annotations of each element with its Kα energy level (C: 0.284, O: 0.532, Na: 1.071, Mg: 1.253, Al: 1.486, P: 2.013, S: 2.307, Cl: 2.621, Ca: 3.69 (k eV)) (B, F). The metal peaks were mapped for the telson (C) and the exoskeleton (G). The composition of elements was calculated from the total spectrum counts (D: 320,357, H: 141,002).

**S3 Figure SEM/EDS analysis of *H. gigas* captured from the Izu-Ogasawara Trench.**

An SEM/EDS analysis was conducted on *H. gigas* captured from the Izu-Ogasawara Trench. Four views of the exoskeleton were analyzed (A, E, I, M) as described in the Methods section. Panels E, I, and M were observed with accelerating voltages of 15 kV, because oil components induce sample charging and cause drift in SEM/EDX images, which cannot be suppressed at high acceleration voltage sufficiently. Only calcium and aluminum were mapped (B, F, J, N). The EDS spectrum includes an annotation of each element with its Kα energy level (C: 0.284, O: 0.532, Na: 1.071, Mg: 1.253, Al: 1.486, P: 2.013, S: 2.307, Cl: 2.621, Ca: 3.69 (k eV)). All EDS signals were detected and calculated from the total spectrum counts (C and D, G and H, K and L, O and P).

**S4 Figure Phylogenetic tree of the captured coastal amphipods and related amphipods reconstructed from the mitochondrial COI protein sequence alignment (192 amino acids).**

The COI genes were amplified and cloned in *E. coli* DH5α as described in the Methods section. Then, we decided DNA sequences of 6 *E. coli* clones. The amino acid sequence of the COI obtained from the coastal amphipods indicated “clone1-6” in this study. The amino acid sequences of the COI of related amphipods were obtained from GenBank, and each accession number was added after the species name. Bold lines indicate bootstrap support above 95% as inferred from the maximum likelihood analysis. The scale bar for the branch length is denoted by the estimated number of amino acid substitutions per site.

**S5 Figure SEM/EDS analysis of the coastal amphipods.**

Three amphipods were freeze dried and then analyzed. The telson (A, C) and foot (E) of *H. gigas* contained aluminum. Panel A and B were observed with accelerating voltages of 10 kV because of sample movement related to the oil component. EDS analysis was conducted as described in the Methods section (B, D, F). The EDS spectrum includes an annotation of each element with its Kα energy level (C: 0.284, O: 0.532, Na: 1.071, Mg: 1.253, P: 2.013, S: 2.307, Cl: 2.621, Ca: 3.69 (k eV)). The peak of Si was obtained from the backfield in F.

**S6 Figure STEM/EDS analysis of pieces of the deep-sea amphipod’s exoskeleton.**

Exoskeletons of the amphipods captured from the Izu-Ogasawara Trench were removed from the individuals, freeze dried, and then scrapped. Bright-field STEM observations were conducted for pieces of the exoskeletons (A, C, E). Characteristic X-rays were collected over 60 s (B, D, F). The Cu or Mo signals were caused by the TEM grid. The Si signal was background.

**S7 Figure *H. gigas* individuals used for the aluminum measurements.**

Deep-sea amphipod *H. gigas* individuals were immediately frozen and maintained at −80°C after capture from Challenger Deep. These amphipods were selected randomaly from frozen stock.

**S8 Figure SEM/EDS analysis of the exoskeleton of the head of *H. gigas*.**

*H. gigas* specimens captured from Challenger Deep were freeze dried for the SEM observations (A). SEM observations were conducted without any coating. The EDS spectrum included an annotation of each element with its Kα energy level (C: 0.284, O: 0.532, Na: 1.071, Mg: 1.253, Al: 1.486, Si: 1.739, P: 2.013, S: 2.307, Cl: 2.621, Ca: 3.69 (k eV)) (B). The total spectrum counts were 141821 in the EDS analysis, and the signals of Si, Al and Ca were mapped (C).

**S9 Figure Effect of gluconic acid/gluconolactone removal on aluminum extraction capacity of *H. gigas* body fluid** Three *H. gigas* individuals were used for the experiment. We prepared *H. gigas* body fluid and removed gluconic acid/gluconolactone from *H. gigas* body fluid with enzymes as described in Materials and Methods (enzyme treated sample. Control sample was prepared without enzyme (control).

**S10 Figure Extraction of iron from the sediment of Challenger Deep.**

The sediment was suspended in 25 mM sodium acetate buffer (pH 5.0) containing 10 mM sodium gluconic acid/gluconolactone or not (control). The suspension was pressurized at 100 MPa and incubated at 2°C for 1 h. After decompression, the sediment was separated with centrifugation (15,000 x g at 4°C for 2 min). The iron content of the supernatant was measured as described in the Methods section. The error bar shows the S.D. (n=3).

**S11 Figure Effect of aluminum hydroxide gel on the release of protein from salmon roe under high pressure.**

Washed salmon roe were soaked and pressurized in artificial seawater at 100 MPa, which is the same pressure observed at approximately 10,000 m in depth, or in the control at 0.1 MPa, which is the same pressure observed in the atmosphere at sea level, for 24 h in a pressure-resistant bottle. The salmon roe that suffered the greatest damage after decompression are displayed in panel A. After decompression, the protein content of the artificial seawater was measured (panel B). The error bar shows the S.D. (n=4). The bar in panel A indicates 1 cm.

Supplementary Table 1 Amount of aluminum in the body of *H. gigas*

Supplementary Table 2 The TEM/EDS analysis of the sediments

Supplementary Table 3 Metabolic analysis of *H. gigas*

Supplementary Table 4 Amount of gluconic acid in the body of *H. gigas*

